# Improving the diagnosis and classification of Ph-negative myeloproliferative neoplasms through deep phenotyping

**DOI:** 10.1101/762013

**Authors:** Korsuk Sirinukunwattana, Alan Aberdeen, Helen Theissen, Nikolaos Sousos, Bethan Psaila, Adam J. Mead, Gareth D.H. Turner, Gabrielle Rees, Jens Rittscher, Daniel Royston

**Affiliations:** Institute of Biomedical Engineering (IBME), Department of Engineering Science, Old Road Campus Research Building, University of Oxford, UK; Big Data Institute / Li Ka Shing Centre for Health Information and Discovery, Old Road Campus, University of Oxford, UK; Oxford NIHR Biomedical Research Centre, Oxford University Hospitals Trust, Oxford, UK; Medical Research Council (MRC) Molecular Haematology Unit, MRC Weatherall Institute of Molecular Medicine, University of Oxford, UK; Haematopoietic Stem Cell Biology Laboratory, Medical Research Council (MRC) Weatherall Institute of Molecular Medicine, University of Oxford, UK; Department of Cellular Pathology, John Radcliffe Hospital, Oxford University NHS Foundation Trust, Oxford, UK; Ludwig Institute for Cancer Research / Nuffield Department of Medicine, Old Road Campus Research Building, University of Oxford, UK; Nuffield Division of Clinical Laboratory Sciences, Radcliffe Department of Medicine, John Radcliffe Hospital, University of Oxford, UK

**Keywords:** pathology, artificial intelligence, myeloproliferative neoplasm, megakaryocyte, bone marrow trephine, image analysis

## Abstract

Myeloproliferative neoplasms (MPNs) are clonal disorders characterized by excessive proliferation of myeloid lineages. Accurate classification and appropriate management of MPNs requires integration of clinical, morphological and genetic findings. Despite major advances in understanding the molecular and genetic basis, morphological assessment of the bone marrow trephine (BMT) remains paramount in differentiating between MPN subtypes and reactive conditions. However, morphological assessment is heavily constrained by a reliance on subjective, qualitative and poorly reproducible criteria. To address this, we have developed a machine-learning strategy for the automated identification and quantitative analysis of megakaryocyte morphology using clinical BMT samples. Using a sample cohort of recently diagnosed or established ET (n = 48) and reactive control cases (n = 42) we demonstrated a high predictive accuracy (AUC = 0.95) of automated tissue ET diagnosis based upon these specific megakaryocyte phenotypes. These separate morphological phenotypes showed evidence of specific genotype associations, which offers promise that an automated cell phenotyping approach may be of clinical diagnostic utility as an adjunct to standard genetic and molecular tests. This has great potential to assist in the routine assessment of newly diagnosed or suspected MPN patients and those undergoing treatment / clinical follow-up. The extraction of quantitative morphological data from BMT sections will also have value in the assessment of new therapeutic strategies directed towards the bone marrow microenvironment and can provide clinicians and researchers with objective, quantitative data without significant demands upon current routine specimen workflows.

## Introduction

Recent advances in computational image analysis have the potential to transform the conventional morphological assessment of human tissues.^1,2^ The quantitative assessment of specific cell populations and the systematic description of tissue architecture have application in replacing or augmenting the subjective categorical classification systems that are central to the diagnosis of many human cancers. Moreover, translation of advanced tissue and single-cell-based genomic and proteomic technologies into new therapeutic strategies will require sophisticated approaches to the assessment of complex pathological tissues that are beyond the scope of routine histopathology.

Philadelphia-negative myeloproliferative neoplasms (Ph-MPN) are driven by acquired mutations in the JAK-STAT signalling pathway of haematopoietic stem cells, resulting in the excessive proliferation of one or more blood lineages.^3^ The three most common Ph-MPNs (essential thrombocythaemia [ET], polycythaemia vera [PV] and primary myelofibrosis [PMF]) have overlapping clinical and laboratory features that can make their distinction challenging, particularly at early disease time points.^4^ In 95% of driver mutation-bearing cases, the MPN phenotype is accounted for by somatic mutations in 3 genes: *JAK2*, *CALR* and *MPL*. Mutations in non-MPN-driver genes also occur in up to one half of MPNs, some of which influence overall, leukaemia-free and myelofibrosis-free survival.^5–7^ Although detection of one or more of these mutations identifies the case in question as clonal and is useful in eliminating a number of reactive differential diagnoses, they are not disease specific.^8^

Approximately a third of ET and PMF cases lack a mutation in one of the three main MPN driver genes (so called triple negative [TN]).^9^ The distinction between TN ET cases and a ‘reactive’ process, for example due to chronic inflammation, remains particularly challenging. This is reflected in the revised 2016 WHO Classification scheme of myeloid malignancies, in which particular emphasis is placed upon the integration of clinical, genetic and histological features for the diagnosis of ET and other MPNs.^10^ Central to the histological interpretation of bone marrow trephines (BMT) from suspected MPN patients is the assessment of megakaryocytes using long-established but subjective descriptions of their cytological and topographic features. These include variations in cell size, atypia (nuclear lobulation / complexity etc.) and cell clustering which may be subconsciously based on assessment of only a subset of the megakaryocytes examined in the tissue section. Despite their continued incorporation into the 2016 WHO classification system, there is controversy about the relative significance and reliability of these subjective cytological descriptions, with several studies documenting considerable intra- and inter-observer variability, even amongst experienced haematopathologists.^11,12^ An improved, quantitative approach to the description / analysis and classification of the complex cytological features of BMT megakaryocytes has the potential to significantly enhance the histological component of integrated MPN diagnosis.

We propose that the complex cytological features of megakaryocytes can be captured on digital images of routinely prepared haematoxylin and eosin (H&E) stained sections, and be used to develop improved disease diagnosis and classification tools through image analysis and artificial intelligence (AI) technologies. These approaches offer a unique opportunity to identify genotype-phenotype correlations that can inform future clinical decision making and basic research into patients with MPNs. Recent advances in computational image analysis have already shown significant promise in various histological diagnosis / classification tasks in common human malignancies.^13–19^ In the latter context, image analysis has even demonstrated the potential to provide robust predictions about key, targetable mutations.^20^

Here we describe a computational method of subtyping megakaryocytes based on their cytomorphological features and determining their relative association with an underlying diagnosis of MPN (ET) or a reactive / non-neoplastic condition. We developed a platform that combines manual annotation tools with support from statistical AI models to assist clinical haematopathologists in the efficient identification of megakaryocytes from routine H&E slides of clinical BMTs. Clustering analysis based on deeply-learned features was deployed to identify megakaryocyte phenotypes and we demonstrate the existence of 11 distinct megakaryocyte subtypes within the marrow of normal and ET BMTs. Our analysis reveals a clear association between the distribution frequency of particular megakaryocyte phenotypic subtypes and a diagnosis of ET. Furthermore, we provide evidence of a direct association between particular megakaryocyte phenotypes and the underlying ET mutational status. Importantly, although the study samples represent a modest cohort of ET (n=48) and reactive control cases (n=42), our analysis was restricted to a single cell population and resulted in a library comprising more than 25,000 megakaryocytes.

In contrast to the high costs associated with specialised sequencing and bioinformatics facilities, the approaches outlined here rely upon H&E stained sections that are prepared as part of the routine investigations of MPN patients. Taken together, these results hold significant promise in the future investigation and clinical management of MPN patients, with the added value of being well suited to integration into existing diagnostic workflows including those operating in low resource settings.

## Materials and Methods

### Clinical Samples

BMT samples were obtained from the archive of the host institution. Whole slide scanned images (Hammamatsu NanoZoomer 2.0HT / 40X / NDPI files) were prepared from 4 μm H&E stained sections cut from formalin fixed paraffin-embedded (FFPE) blocks. The data set comprised 90 samples (48 ET and 42 reactive / non-neoplastic) with reactive cases derived from a range of patients undergoing lymphoma or plasma cell myeloma investigation or staging in whom there was no evidence of bone marrow involvement and in whom there was no known myeloid disorder. We identified BMTs from the laboratory reporting system or multidisciplinary team meeting (MDT) records. This work was conducted as part of the INForMeD study (Investigating the genetic and cellular basis of sporadic and Familial Myeloid Disorders; IRAS ID: 199833; REC reference: 16/LO/1376; PI: Prof AJ Mead). ET cases represent patients in whom this was either an established or new diagnosis satisfying the diagnostic criteria of the latest WHO classification (2016)^10^. A summary of the key patient characteristics is provided in Supplemental Table 1.

### Constructing a Megakaryocyte Library — a Human-in-the-loop Approach

Supervised machine learning methods are dependent on large numbers of labelled examples. The analysis of megakaryocyte morphology presented here required close collaboration with specialist haematopathologists in order to ensure that a reliable, high-quality and well-curated dataset (library) contained suitable example images of megakaryocytes. These example images included megakaryocytes with the wide variety of cytomorphological features commonly encountered in normal and diseased BMT samples.

In an attempt to reduce the burden of repetitive and labour-intensive manual annotation, we designed assistive AI models to enable quick and efficient feedback from their annotation tasks. For this we used a human-in-the-loop (HITL) iterative method that leverages both human and machine judgement. The loop refers to the cycle in which humans provide feedback to tune the latest model; the latest model updates the predictions and further feedback is provided.

### Iterative Pipeline

To begin with, we placed all BMT slide images in the unlabelled pool (Figure 1a). To train our initial machine learning algorithms for identification (Figure 1a) and delineation (Figure 1b) of megakaryocytes, we created a first limited set of training data. A random selection of 14 reactive and 6 ET slide images were annotated manually by a specialist haematopathologist (DR). The annotation process required delineating the boundary of every megakaryocyte in each image. This direct manual annotation ensured that the number of examples in the dataset was above the threshold required to train the first iteration of the detection and segmentation models (Figure 1a-b).

**Figure 1.**
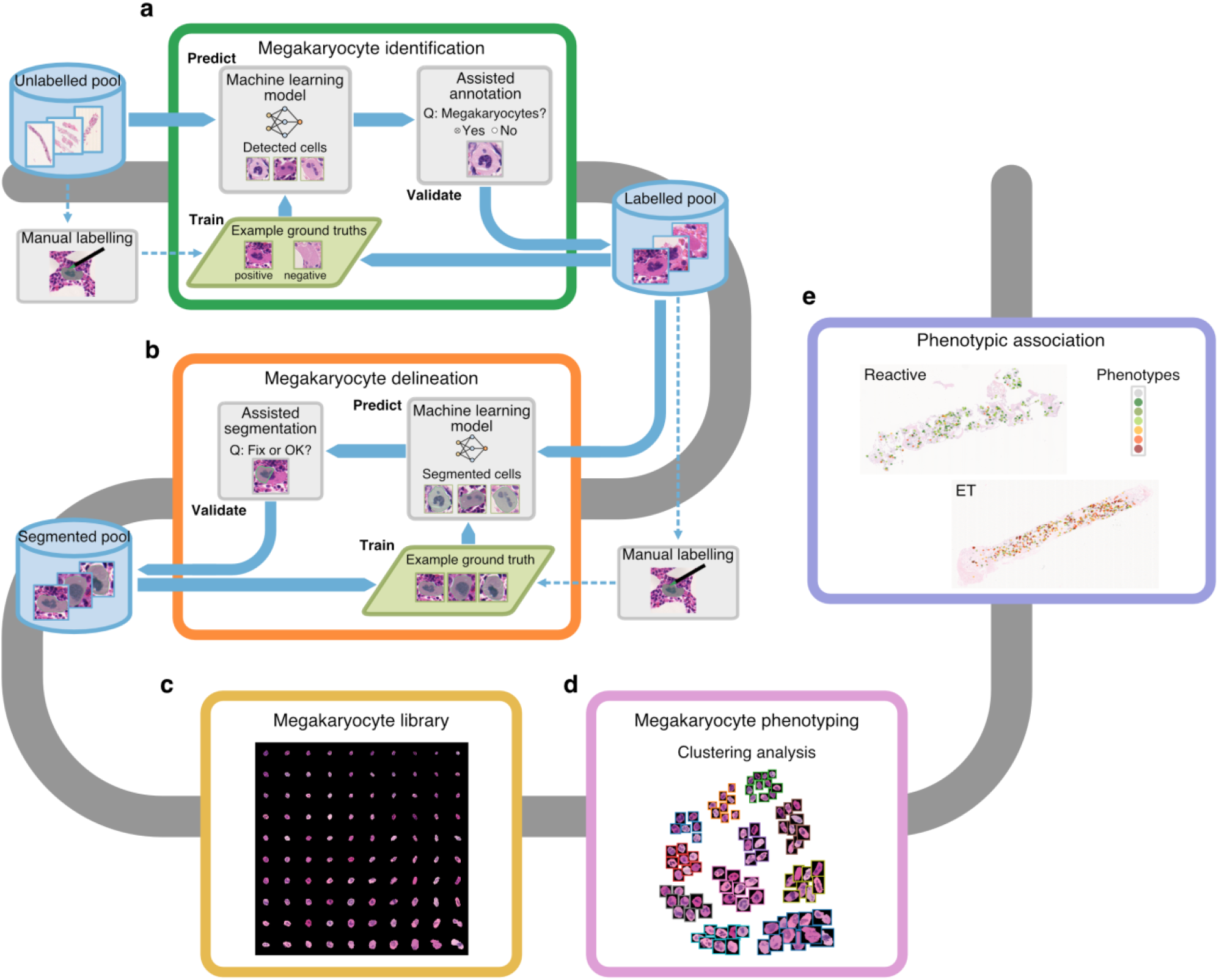
Overview of the Computational Pipeline for Phenotypic Analysis of Megakaryocytes. In order to effectively build an annotated library of megakaryocytes, assisted annotation tools for identification (**a**) and delineation (**b**) have been developed. A library containing at least 25,000 annotated megakaryocytes from samples with diverse disease background could be generated within a short amount of time (**c**). Clustering analysis performed on the library of megakaryocytes to identify possible phenotypes (**d**). Statistical analyses were carried out to assess the diagnostic associations of different megakaryocytes phenotypes (**e**).

After the first detection and segmentation models were trained, the annotation paradigms were switched to the iterative HITL feedback approach. This system consists of iterating through three distinct steps (Figure 1a-b):

1. **Train:** The validated annotation cases were randomly allocated to either the training or evaluation datasets, which accumulated validated examples from all iterations. The models were trained on the training set, with performance measured on the evaluation set.
2. **Predict:** The updated models were used to detect and delineate megakaryocytes in a sampled subset of unseen cases.
3. **Validate:** Three specialist haematopathologists (DR, GR and GT) provided feedback on the inference results (predicted megakaryocytes for the prediction model and predicted boundaries for the segmentation model), indicating whether the inference results were valid. Validated results were then placed in the labelled pool for detection and segmented pool for segmentation (Figure 1a-b).

For the complete details of all iterations of the training and validation procedure, see Supplemental Tables 2 and 3.

### Detection

The detection task required predicting the location and dimensions of a rectangular box bounding each megakaryocyte on a BMT slide (Figure 2a). We used a deep neural network called Single Shot Multibox Detector.^21^ This method defines a default set of bounding boxes over different aspect ratios and scales. To find the megakaryocytes, the network generated a score for each default box to indicate the likelihood that it contained a megakaryocyte and a score for the recommended offset for each default box that more closely matches the identified megakaryocyte. For complete details of the training method see Supplemental Methods. The validity of each predicted bounding box was confirmed by at least one specialist haematopathologist. To collect the specialist feedback as quickly as possible, we simplified the interaction in our custom web interface (Figure 2a). The interface cycled through the predictions and the user simply assigned a ‘positive’ or ‘negative’ label to each prediction with the labels mapped to scores of +1 and −1, respectively. The level of consensus was quantified by showing the same example megakaryocyte to more than one pathologist. If the average score was positive, the candidate cell was labelled as a megakaryocyte.

**Figure 2.**
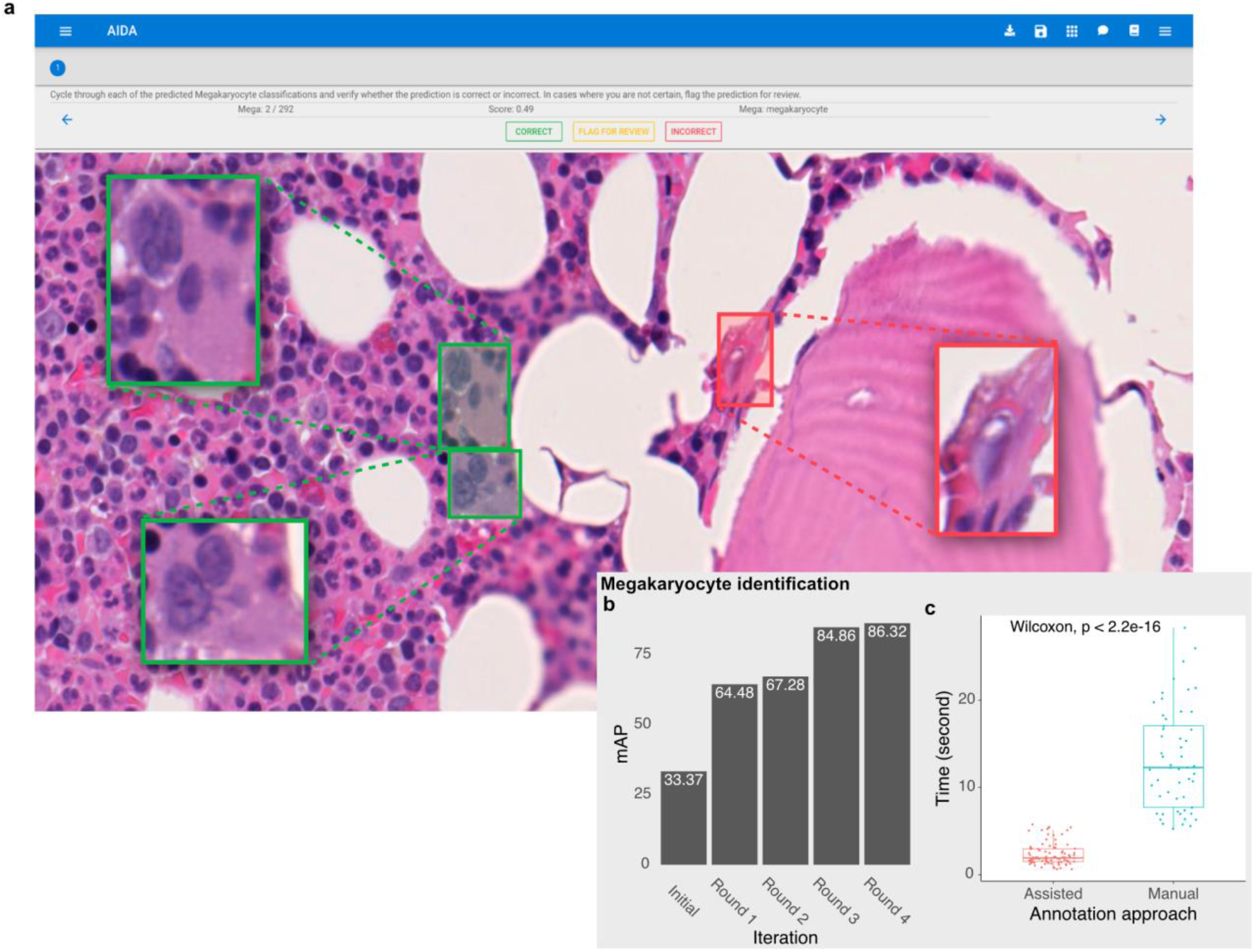
Efficiency of the Assisted Tool for Megakaryocyte Identification. Our web-based interface for assisted annotation. Candidate megakaryocytes are automatically identified by the AI algorithm, allowing haematopathologists to quickly review and confirm these as megakaryocytes (green boxes) or non-megakaryocytes (red boxes) (**a**). The human-in-the-loop assisted annotation tool could achieve a high level of accuracy as measured by the mean average precision (mAP) within a few training iterations (**b**), and the amount of annotation time is significantly reduced by using our tool (**c**). Wilcoxon rank-sum test, significance level = 0.05.

### Segmentation

Image segmentation involves partitioning an image into different regions by assigning a class label to each pixel such that pixels of the same class share similar characteristics. Segmentation is often used to locate the boundaries of objects of interest; in this case megakaryocyte cells. Supplemental Figure 1a shows an example of a segmented megakaryocyte. We used a convolutional neural network U-Net to delineate the boundaries of megakaryocytes.^22^ Our implementation of the network produced both a segmentation result and an estimated score for the quality of the result. The estimated intersection over union (IoU) score, ranging between 0 and 1, reflected how well the segmentation result overlapped with the ground truth. For complete details of the training of the method see Supplemental Methods. In each iteration of the iterative training cycle, the model was applied to unseen data. We then selected segmentation results that are considered outliers based on Tukey's rule (the estimated IoU < Q1 − 1.5 [Q3 − Q1], where Q1 and Q3 denote the 1st and 2nd quartile of the estimated IoU values of segmentation results). These selected segmentation results deemed to be outliers went through a quality control stage involving inspection and correction by two annotators (KS and HT) [Figure 1b].

### Megakaryocyte Phenotyping

We used two methods to identify morphological phenotypes: feature extraction and clustering analysis. For feature extraction, it was important to make sure that only the morphology of megakaryocytes impacted the resulting output. To account for this, we used the fully segmented megakaryocyte image patches (see Section Segmentation) and removed the background area from the image. We used the autoencoder method to transform the image patches into numerical vectors (latent representations) of length 128.^23^ Each vector is an efficient encoding of the megakaryocyte morphology. For complete details of the training method see Supplemental Methods.

Following feature extraction, we performed a clustering analysis to group morphologically similar megakaryocytes. We trained a self-organizing map (SOM) on the learned latent representation vectors to produce a 10-by-10 grid with 100 groups of similar megakaryocytes.^24^ To further reduce the number of clusters we applied Markov clustering on the self-organising groups.^25^ We chose the Markov clustering result that maximizes the modularity measure.^26^ The results from Markov clustering and self-organising map were used to assign each megakaryocyte to a cluster (Figure 1d; Figure 3a).

**Figure 3.**
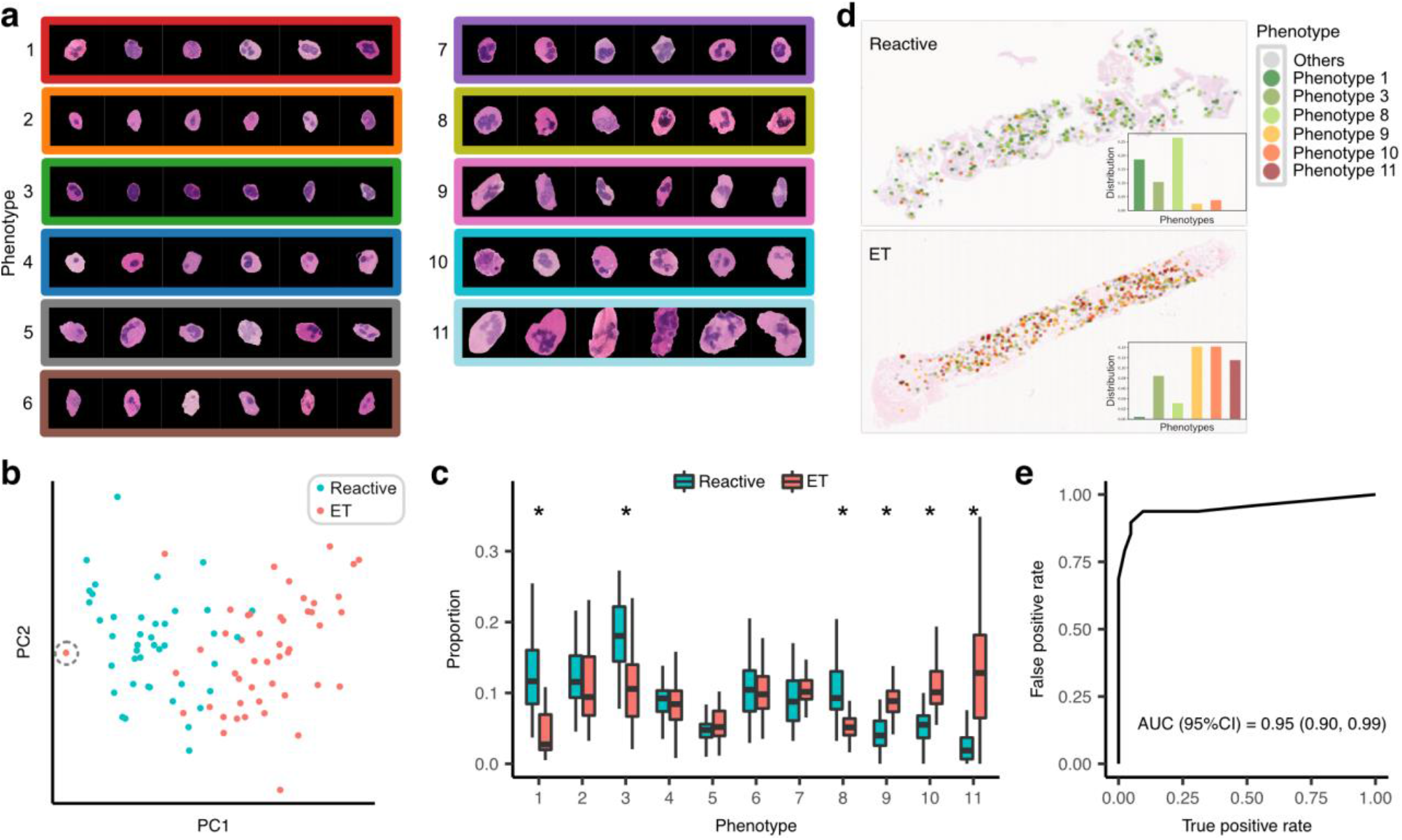
Statistical Analyses of Megakaryocyte Phenotypes. A total of 11 megakaryocyte phenotypes were automatically discovered in the unsupervised clustering analysis (**a**). Proportions of 11 megakaryocyte phenotypes could be used to describe a BMT sample. The results from principal component analysis at the sample level show that reactive and ET subgroups are reasonably separated under the identified phenotypic profiles (**b**). Association analysis of the phenotypic distributions and the diagnostic status (Bonferroni adjusted Wilcoxon rank-sum test, statistical significance at 0.05). An asterisk indicates a statistically significant result. [A dashed circle indicates the outlier sample referred to in the results section]. (**c**). Detailed spatial distribution of different megakaryocyte subtypes on examples of a reactive and ET bone marrow trephine. Megakaryocytes of phenotypes 1,3, and 8 are prominent in the reactive sample, while phenotypes 9-11 are most prominent in the ET sample (**d**). The k-nearest neighbour classifier (k = 9) reached the AUC of 0.95 demonstrating the potential to accurately discriminate ET from reactive samples based on megakaryocyte phenotypes (**e**).

### Spatial Distribution Analysis

We estimated the spatial density of megakaryocytes using kernel density with an Epanechnikov kernel. A single kernel’s bandwidth was estimated for all samples by Silverman’s rule-of-thumb.^27^ Clusters of densely packed megakaryocytes have uniformly high values of density and so low variance of the density within clusters. We selected a global cut-off that yields the lowest average variance of density within clusters to determine clusters of densely packed megakaryocytes. Figure 4a shows examples of megakaryocyte clusters.

**Figure 4.**
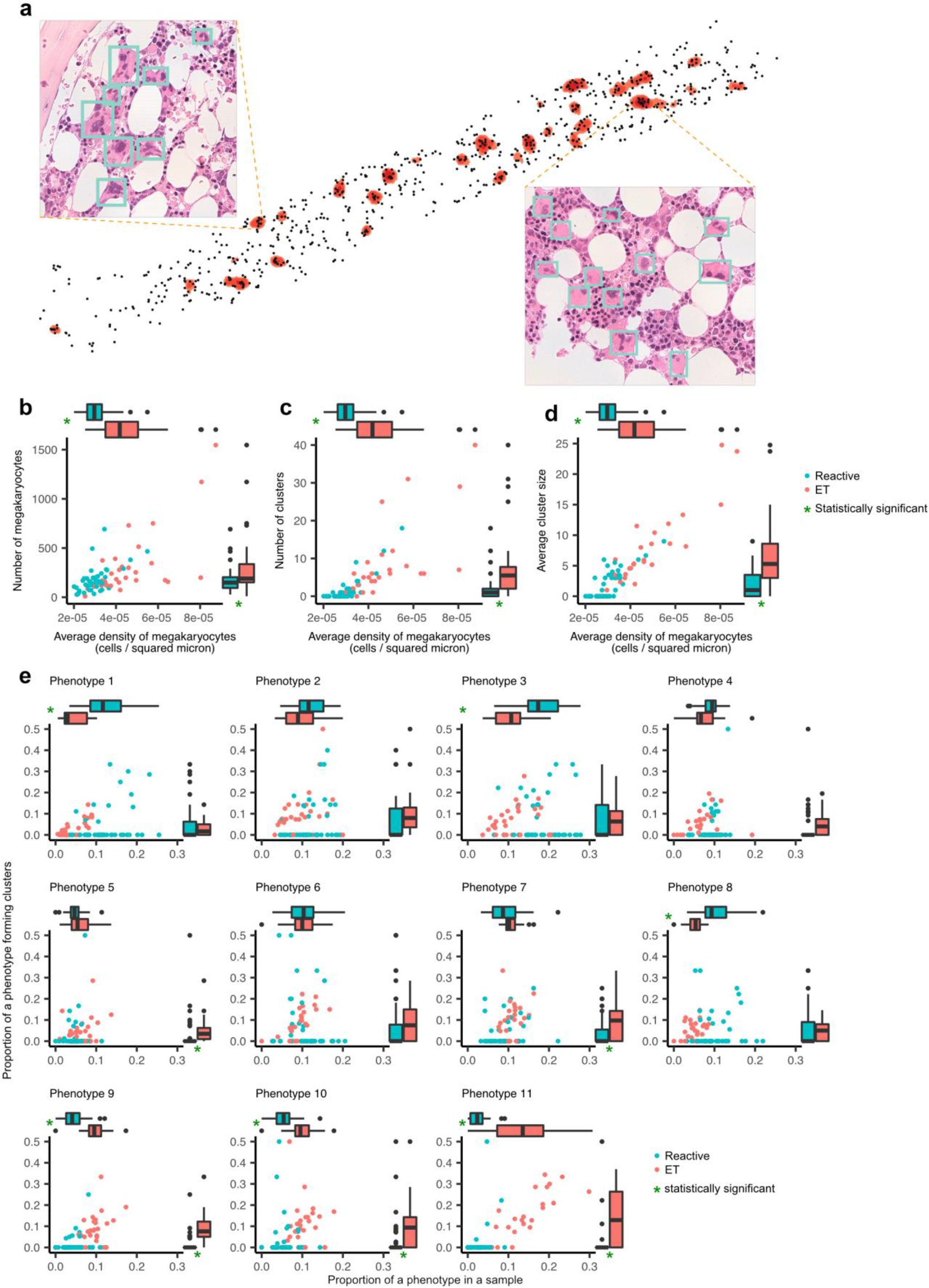
Spatial Statistics of Megakaryocytes. Clusters of densely packed megakaryocytes (orange) could be identified by our analytical pipeline (**a**). The average density of megakaryocytes in ET is significantly higher than that of the reactive samples (Wilcoxon rank-sum test, p < 0.05) [**b-d**] and there is a strong correlation between average density and the number of megakaryocytes on a sample (**b**), the number of clusters (**c**), and the average cluster size (**d**). The number of cells, cluster size and number of clusters is significantly higher in ET compared to reactive samples (Wilcoxon rank-sum test, p < 0.05). Although phenotypes 1,3, and 8 are significantly enriched in reactive samples (Bonferroni adjusted Wilcoxon rank-sum test, p < 0.05), they are not enriched in clusters of densely packed megakaryocytes. By contrast, phenotypes 9-11 are significantly enriched in ET samples and tend to form clusters of densely packed cells (Bonferroni adjusted Wilcoxon rank-sum test, p < 0.05) (**e**).

### Case Classification

A *k*-nearest neighbour classifier utilising six discriminative megakaryocyte phenotypes (Phenotypes 1, 3, 8, 9, 10, and 11) [Figure 3c] was trained to discriminate reactive and ET samples. The distance between samples is defined by Euclidean distance. We used 5-fold cross-validation in which each cross-validation fold preserves the ratio of reactive to ET cases to maintain the class balance at their original levels. Classification accuracy measures including precision, recall, F1-score, and area under the receiver operating characteristic curve (AUC) were recorded for each cross-validation permutation. To evaluate the effect of the number of nearest neighbours, *k*, on the classification performance, we trained separate classifiers using different values of *k* (*k* = 1,3,5,7, and 9). See Figure 3e and Supplemental Table 7 for the classification results.

### Implementation

All machine learning models were implemented using Python and the deep learning Pytorch library.^28^ Statistical analyses were performed using R version 3.5.1. Our web-based annotation tool is publicly available at https://github.com/alanaberdeen/AIDA.

## Results

### Human-in-the-loop Assisted Annotation Tools to Accelerate the Curation of Megakaryocytes

We employed a human-in-the-loop methodology to efficiently build a large library of annotated megakaryocytes (approx. 25,000 megakaryocytes). Figure 2a shows the interface of our web-based annotation tool for megakaryocyte identification. The identification tool detected candidate megakaryocytes for which the delineation tool suggested segmentation masks (Figure 1b; Supplemental Figure 1a). To ensure accuracy and quality, these predicted results were reviewed and edited by haematopathologists and fed into the AI models for further training to iteratively improve the model performance (Figure 2b; Supplemental 1b-c). Our tools achieved high levels of accuracy for identification (mAP = 0.86) and delineation (IoU = 0.93) within four training iterations. The time spent on annotating megakaryocytes with our assisted AI tools was significantly less than manual annotation (Figure 2c; Supplemental Figure 1d).

### Megakaryocyte Phenotypes and Their Diagnostic Associations

To identify a feature set that best captures megakaryocyte morphology an autoencoder neural network was used. A total of 11 morphological subtypes were identified through clustering analysis performed on these learnt features. Certain of the 11 identified phenotypes have distinct, readily appreciated morphological characteristics (Figure 3a; Supplemental Figure 2). For instance, cells identified as belonging to phenotype 11 are significantly enlarged and frequently contain an atypical, polylobated nucleus. By contrast, cells of phenotype 3 are relatively small with a high nuclear-cytoplasmic ratio. However, certain other megakaryocyte phenotypes are not so easily distinguished by trained haematopathologists, which emphasizes the limitations of conventional subjective assessment. As predicted, analysis of megakaryocyte spatial distribution revealed that ET samples contain an increased cell density and absolute cell number when compared to reactive samples (Figure 4a-b), with megakaryocytes in ET tending to form increased numbers of large, more densely packed clusters (Figure 4c-d).

We used the proportion of megakaryocytes of each phenotype to characterise the BMT samples. Principal component analysis demonstrated separation between reactive and ET samples under the identified phenotypic profiles (Figure 3b). We identified significant enrichment of phenotypes 1,3, and 8 in reactive samples and phenotypes 9-11 in ET samples (Figure 3c). Detailed visualisation of the spatial distribution of megakaryocytes on the scanned images confirmed the prominence of these phenotypes in reactive and ET samples (Figure 3d). Although significantly enriched in reactive samples, the clustering of megakaryocytes of phenotypes 1, 3, and 8 was not different between reactive and ET samples. By contrast, phenotypes 9-11 were both significantly increased and more likely to form clusters in ET (Figure 4e). Based on the proportion of these discriminative phenotypes, we trained *k*-nearest classifiers with different values of *k* to classify ET (n = 48) and reactive samples (n = 42). The optimal classifier used *k* = 9 and reached the AUC of 0.95 in 5-fold cross validation (Figure 3e; Supplemental Table 7).

### Morpho-molecular Association Analysis

We performed statistical analysis to assess the association between megakaryocyte phenotypes and the underlying mutational status, including TN cases and those carrying the two most common driver mutations (*JAK*2 and *CALR*). Statistically significant associations to the *CALR* mutational status were observed for megakaryocyte phenotypes 4 and 7 (Figure 5a). Indeed, principal component analysis also showed a reasonable separation of *CALR* mutated samples from TN and *JAK2* mutated samples (Figure 5c). No significant morpho-molecular associations were identified for cases carrying a *JAK2* mutation or having TN status (Supplemental Figure 3). The proportion of megakaryocytes with phenotype 11 was significantly higher in ET samples than reactive, and there was an increasing trend in the proportion of phenotype 11 amongst the ET samples as stratified by order of TN, *JAK2*, and *CALR* mutation status (Figure 5b). Furthermore, we observed a moderate correlation between the proportion of phenotype 11 cells and the peripheral blood platelet counts (Spearman correlation coefficient = 0.43) [Figure 5d-e].

**Figure 5.**
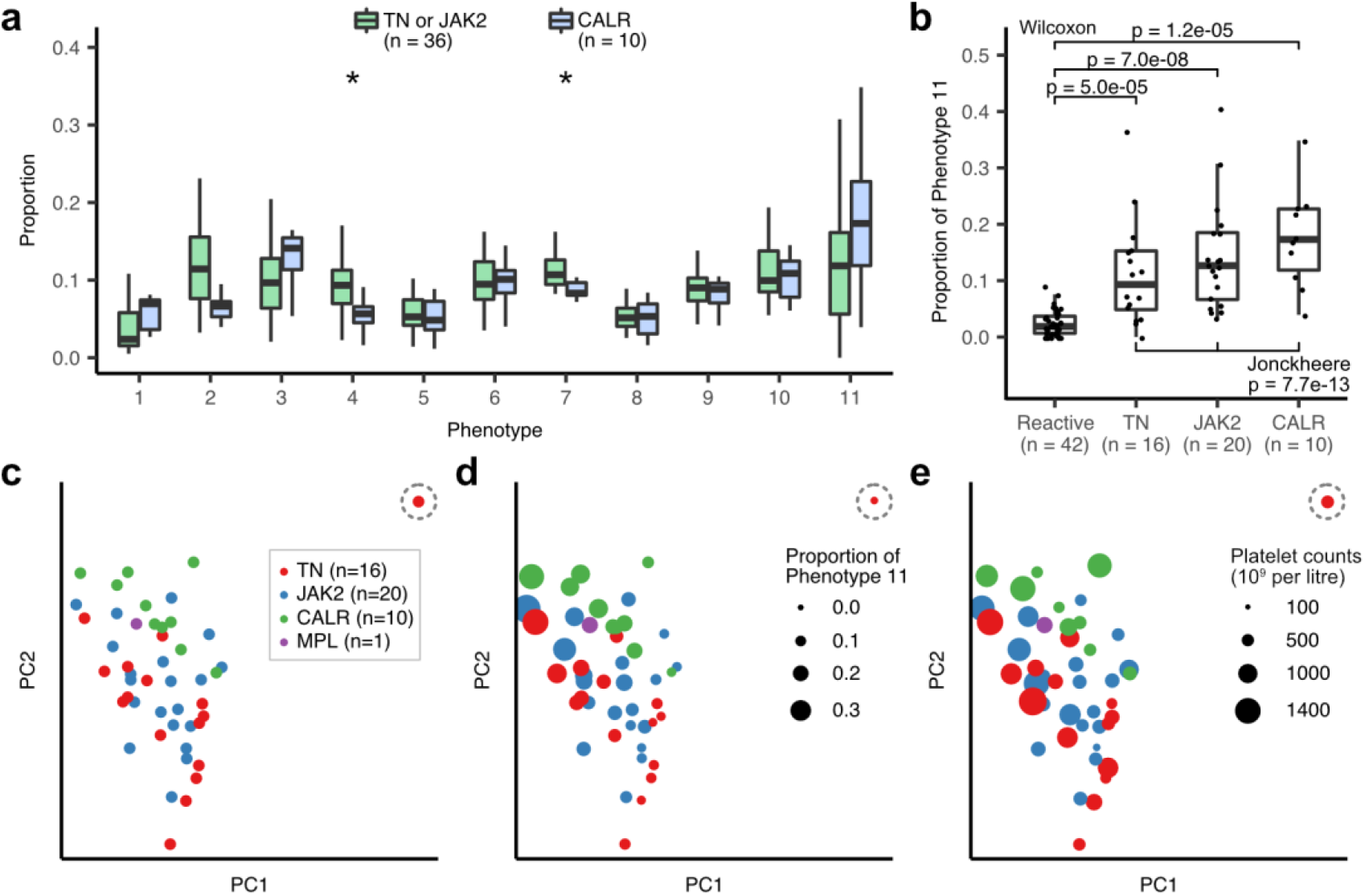
Morpho-molecular Associations in ET. Associations between phenotypes and mutational status in ET samples (Bonferroni adjusted Wilcoxon rank-sum test, significance level = 0.05) (**a**). Associations between the mutational status and the proportion of the phenotype 11 (Bonferroni adjusted Wilcoxon rank-sum test, significance level = 0.05). Trend in phenotype 11 amongst ET samples (as stratified in the order of TN, *JAK2* and *CALR* mutation status) was tested using Jonckheere’s test (significance level = 0.05) (**b**). Principal component analysis of the diagnosed samples based on the phenotypic profiles (**c**), variation in the proportion of phenotype 11 (**d**) and the platelet count (**e**). A dashed circle indicates the outlier sample referred to in the results section.

Representative samples from our analysis are illustrated in Figure 6. A typical reactive sample (patient 1) can be readily distinguished from those of ET patients with or without an identified driver mutation (patient 7 representing the only *MPL* mutant case in our cohort of 48 ET samples). Of interest, the sample from patient 4 harbouring a *JAK2* mutation with a low variant allele frequency (VAF <5%) was identified by our analysis as being phenotypically more similar to the TN samples. Notably, while clearly identified as cases of ET using our current analysis, the samples from patients 2 and 4 were challenging cases to interpret morphologically. Indeed, in both cases the original histology reports offered a descriptive account of the specimen and specifically raised a reactive differential diagnosis. This likely reflects the relatively low proportion of megakaryocytes with phenotypes 10 and 11, which are more readily appreciated on routine, subjective assessment. Finally, the BMT specimen from patient 8 was processed and analysed as a case of ET but is clearly identified as an outlier in Figure 3b and Figure 5c-e. Subsequent review revealed this case to have been incorrectly assigned (typographical error) as ET rather than reactive during image acquisition and labelling prior to our analysis. This highlights the discriminatory potential of our analytical tools.

**Figure 6.**
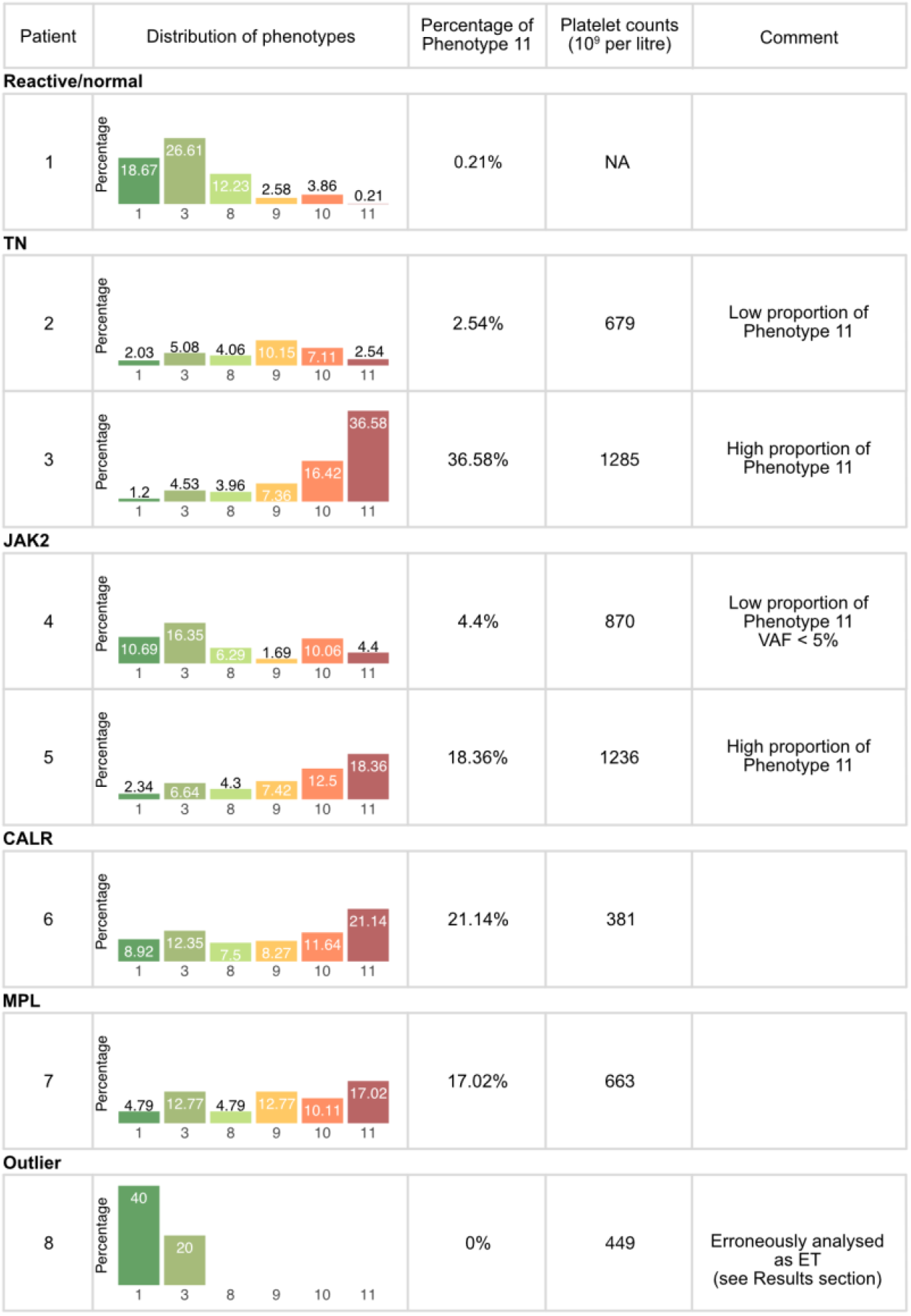
Representative examples of reactive, ET, and ‘outlier’ BMT samples.

## Discussion

The BMT histological criteria required for an MPN diagnosis based on current classification schemes are qualitative and rely upon the subjective assessment of morphological features by a trained haematopathologist.^10^ In particular, there is emphasis on the description of various cytological and topographical features of megakaryocytes, with various accounts of cellular atypia / pleomorphism, nuclear lobulation and clustering outlined in most diagnostic manuals and textbooks. These subjective, qualitative descriptions are poorly reproducible, even between experienced haematopathologists.^11,12^

Here we present a strategy for the annotation, extraction and quantitative description of MPN megakaryocytes using advanced machine learning approaches. The automated capture of cellular features and their translation into reproducible unsupervised morphological phenotypes allows the complex range of megakaryocytes to be quickly and efficiently curated from routine BMT samples, and overcomes the limitations of describing this complexity in language that is subject to interpretation. These approaches have identified specific megakaryocyte phenotypes that offer significant promise as automated adjuncts in the diagnosis of MPN and their distinction from potential reactive mimics. In our cohort of ET and reactive / normal BMT samples, principal component analysis using the identified megakaryocyte phenotypes showed a clear separation of these cases, with 6 out of 11 specific phenotypes being significantly associated with either a reactive or ET diagnosis. As expected, several of these phenotypic groups broadly correspond to the traditional phenotypes described in conventional subjective classification schemes. However, our analysis suggests that particular phenotypic groups are not easily distinguished by conventional approaches and some may have significance beyond the simple segregation of neoplastic and non-neoplastic cases. Specifically, megakaryocyte phenotypes 4 and 7 appear to be disproportionately represented in TN and *JAK2* cases of ET when compared to *CALR* mutation-bearing samples, although interestingly these phenotypes were not identified as being significantly over-represented in the ET cohort as a whole. Taken together, these observations suggest that in addition to having utility in the distinction of ET from reactive mimics, particular megakaryocyte phenotypes correspond to specific genetic subgroups of ET. While this finding is supported by recent observations of certain mutational-phenotypic associations in ET ^29^, the identification and characterization of the specific phenotypes presented here could not have been achieved by conventional subjective specimen assessment, and are reliant upon the quantitative data extracted by our automated machine learning tools.

In addition to the identification of 11 distinct megakaryocyte phenotypic groups, our analytical tools allow us to correlate these phenotypes with their topographical distribution within the analysed BMT specimens. As expected, prominent clustering of megakaryocytes was observed in ET, but our analysis also suggests that this clustering of megakaryocytes is associated with particular phenotypes. This observation raises important questions about possible crosstalk between specific clonally related megakaryocyte subpopulations within the bone marrow microenvironment, particularly given the observed morpho-molecular associations, and clearly warrants further investigation.

We consider that the machine learning approaches presented here offer significant promise in several distinct clinical scenarios. Firstly, a fully automated annotation and analysis pipeline of BMT megakaryocytes in suspected cases of MPN has the potential to provide a fast and reliable initial diagnostic assessment of specimens in advance of formal pathology reporting. This is likely to be of value when access to haematopathology expertise is restricted (for example in low-resource healthcare systems). In the context of histological reporting by specialist haematopathologists, we also consider that the automated output from our analysis will provide valuable ancillary information in the routine work up and reporting of BMT samples. In particular, a comprehensive, reliable and easily interpreted summary of the megakaryocytic population will allow the diagnostic pathologist to concentrate on the ‘higher-level’ process of integrating the broader pathological features with the clinical and laboratory findings.^30^ We anticipate that this may prove particularly useful for the assessment of sequential specimens from patients undergoing treatment and / or repeated investigation, in whom quantitative morphological correlates of disease response are currently unavailable.

Importantly, although the morphological assessment of megakaryocytes is central to the histological assessment of MPNs, other features such as marrow cellularity, presence / absence of fibrosis and blast cell estimation are included in the current WHO classification system of MPNs. It is therefore likely that the optimal clinical application of our machine learning megakaryocyte analysis will require structured integration into a more comprehensive quantitative description of the bone marrow microenvironment.

One limitation of our current analysis is the modest number of samples and lack of detailed clinical follow up and molecular characterisation beyond the identification of the key driver mutations *JAK2*, *CALR* and *MPL*. We also recognise that our current study has been restricted to MPN cases labelled as ET and does not include cases of PV or PMF. This partly reflects our intention to establish an analytical workflow with samples in which the morphological analysis of megakaryocytes is not significant disrupted by bone marrow fibrosis, but also recognises that one of the key unmet clinical needs in the management of MPN patients is the challenging distinction of ET and prefibrotic/early PMF (pre-MF). We reason that the machine learning approaches described here will be readily applicable to the interrogation of sample cohorts enriched for cases of pre-PMF and ‘high-risk’ ET, in whom the distinction from low-risk ET is of significant prognostic and therapeutic importance. This will require access to large multi-centre trial cohorts in whom detailed longitudinal clinical and genetic data are available, and is the subject of ongoing work.

Notwithstanding the promise of the machine learning approaches outlined here, important questions remain about the relationship between various molecular / genetic subtypes of MPN and the megakaryocyte phenotypes and their topographical distribution throughout the marrow. Importantly, the molecular landscape of aberrant megakaryopoiesis emerging from novel technologies such as high-throughput single-cell transcriptome profiling is already demonstrating the potential to identify clone-specific cell surface antigens. Similar, modified approaches designed to interrogate intact bone marrow tissue and characterise megakaryocyte sub-populations and their unique microenvironment are clearly required to expand and refine the preliminary findings presented here.

## Acknowledgements

The research was funded by the National Institute for Health Research (NIHR) Oxford Biomedical Research Centre (BRC). JR is supported through the EPSRC funded Seebibyte programme (EP/M013774/1). Computation used the Oxford Biomedical Research Computing (BMRC) facility, a joint development between the Wellcome Centre for Human Genetics and the Big Data Institute supported by Health Data Research UK and the NIHR Oxford Biomedical Research Centre. The views expressed are those of the authors and not necessarily those of the NHS, the NIHR or the Department of Health. HT is funded by the Oxford-Nottingham EPSRC and MRC Centre for Doctoral Training in Biomedical Imaging.

## Authorship Contributions

DR and JR conceived and supervised the study with input from BP and AJM. KS, AA, DR and JR designed the study. AA and KS designed human-in-the-loop protocol for annotation; AA implemented web annotation tools, collected and summarised expert annotations; KS implemented machine learning algorithms. DR, GT and GR verified AI predictions for megakaryocyte identification; KS and HT reviewed AI predictions for megakaryocyte delineation. KS and HT performed statistical analyses. NS, AM and BP identified samples and collected / provided access to clinical and genetic data. KS, AA, HT, JR and DR drafted the manuscript. All authors read and have given approval of the final manuscript.

## Supplementary Material

For further details of the megakaryocyte detection training, segmentation and feature extraction methods please see Supplementary Materials. (Supplementary Figures 1-3, referred to in the main text, are also included in this section).

