## supplemental for "Improving the diagnosis and classification of Ph-negative myeloproliferative neoplasms through deep phenotyping"

### Training of the Megakaryocyte Detection Method

**Input:** For input to the model, the whole-slide images were split into smaller tiles of 512x512 pixels at 40x magnification. (The NDPI files were converted to the tiled pyramid DeepZoom format in order to display the images and also to train on the underlying tiles that were converted to JPEG files.)

**Pretraining:** Initially, there were a limited number of example megakaryocytes in the training set. To support the learning task, the model was first trained on a large set of natural images from the VOC PASCAL dataset.[1] This allowed the low-level knowledge learned in the related task of detecting and classifying natural images to be transferred to the target task of detecting cells in microscopy images. This helped to ensure that the algorithm converged more quickly.

**Data augmentation:** To reduce the chance of overfitting to potential idiosyncrasies, we used extensive data augmentation. This involved applying different transformations to the training data such as flips, scale changes, affine transformations, and colour perturbation to increase the variation in the data.

**Optimal model selection:** During training, the performance of the algorithm was evaluated over time on the hold out 20% of the training data to determine the point of overfitting. The state of the model that yields the maximum detection accuracy was then selected for the inference.

**Counteraction of prediction bias:** In this iterative process, there was a chance that the model would begin to favour a specific subtype of cell. This subtype could have started to dominate the training data and negatively affect the performance of the detection model. To counteract this, we also included 20% of the low confidence (0.4 - 0.6) predictions as well as all predictions with a confidence score > 0.6 in the set of predictions for expert validation. Furthermore, for each iteration of the iterative cycle, we sampled the most recent set of verified megakaryocytes 4 times more frequently than those verified in the previous rounds. As the model has already been trained on previous examples we had more confidence in the predictions. Therefore, we wanted to ensure it was sensitive to differences in the newly verified samples.

**Hyperparameter setting:** For complete details of the hyperparameter values of the model see Supplemental Table 4.

### Training of the Megakaryocyte Segmentation Method

**Input:** Similar to the detection stage, we constructed smaller image tiles for input to the model. The patches were 384x384 pixels at 40x magnification with the identified megakaryocyte at the centre (Supplemental Figure 1a).

**Data augmentation:** During training, we used aggressive spatial and optical augmentations of the data to reduce overfitting.[2]

**Optimal model selection:** We evaluated the performance over time on the hold out 20% of the training data to determine the point of overfitting. The state of the model that yields the maximum segmentation accuracy measured in terms of intersection over union (IoU) was selected for the inference.

**Hyperparameter setting:** See Supplemental Table 5 for details of parameter values of the algorithm.

### Training of Feature Extraction Model

**Input:** We used downsampled patches of size 64x64 pixels. There was also the possibility of an additional confounding effect from non-uniform staining across slides. However, we found that after converting the images to *Lab* colour space the *L* channel, representing pixel lightness, sufficiently encoded the morphological information.[3,4] We could therefore account for non-uniform staining by only using the L channel in further downstream analysis. For training, we used segmented megakaryocytes from all slides.

**Data augmentation:** No further augmentation was applied while training.

**Hyperparameter settings:** For full details on training parameters, see Supplemental Table 6.

**Supplemental Table 1. Clinicopathological Summary of the Cohort**

| | | ET |
|---|---|---|
| | | n = 48 |
| Clinico-pathological features | | |
| Platelet counts (10^9 / L) | | n = 48 |
| | Median (range) | 574 (121, 1384) |
| White blood count (cells / L) | | n = 47 |
| | Median (range) | 7.3 (3.86, 30.9) |
| Haemoglobin (g / L) | | n = 48 |
| | Median (range) | 136 (83, 170) |
| Driver mutations | | |
| TN | | n = 47 |
| | No | 31 |
| | Yes | 16 |
| JAK2 | | n = 47 |
| | Wild-type | 27 |
| | Mutated (V617F) | 20 |
| CALR | | n = 47 |
| | Wild-type | 37 |
| | Mutated | 10 |
| MPL | | n = 47 |
| | Wild-type | 48 |
| | Mutated | 1 |

**Supplemental Table 2. Training and Validation Schedule for the Assisted Megakaryocyte Identification Tool**

| Iteration | Labelled pool | | Unlabelled pool | | Training | | | Validation | | |
|---|---|---|---|---|---|---|---|---|---|---|
| | Reactive | ET | Reactive | ET | Reactive | ET | Mega | Reactive | ET | Mega |
| Initial training | 14 | 6 | 28 | 24 | 7 | 3 | 1069 | 7 | 3 | 1358 |
| Round 1 | 18 | 8 | 24 | 22 | 10 | 4 | 1977 | 8 | 4 | 2411 |
| Round 2 | 22 | 10 | 20 | 20 | 13 | 5 | 2951 | 9 | 5 | 2776 |
| Round 3 | 42 | 13 | 0 | 17 | 27 | 7 | 6904 | 15 | 6 | 4871 |
| Round 4 | 42 | 30 | 0 | 0 | 27 | 19 | 10103 | 15 | 11 | 5947 |

mAP: mean average precision; Mega: megakaryocytes

**Supplemental Table 3. Training and Validation Schedule for the Assisted Megakaryocyte Delineation Tool**

| Iteration | Labelled pool | | Unlabelled pool | | Training | | | Validation | | |
|---|---|---|---|---|---|---|---|---|---|---|
| | Reactive | ET | Reactive | ET | Reactive | ET | Mega | Reactive | ET | Mega |
| Initial training | 14 | 6 | 28 | 24 | 7 | 3 | 1069 | 7 | 3 | 1358 |
| Round 1 | 18 | 8 | 24 | 22 | 10 | 4 | 1977 | 8 | 4 | 2411 |
| Round 2 | 22 | 10 | 20 | 20 | 13 | 5 | 2951 | 9 | 5 | 2776 |
| Round 3 | 42 | 13 | 0 | 17 | 27 | 7 | 6904 | 15 | 6 | 4871 |
| Round 4 | 42 | 30 | 0 | 0 | 27 | 19 | 10103 | 15 | 11 | 5947 |

IoU: intersection over union; Mega: megakaryocytes

**Supplemental Table 4. SSD Parameters**

| Parameter | Value |
|---|---|
| batch size | 16 |
| learning rate | 0.0001 |
| optimizer | SGD |
| - momentum | 0.9 |
| - weight decay | 5.00E-04 |
| - learning rate decay | 9.00E-01 |
| loss function | SSD loss |

SSD: single shot multibox detector; SGD stochastic gradient descent

**Supplemental Table 5. UNet Parameters**

| Parameter | Value |
|---|---|
| batch size | 32 |
| learning rate | 0.0002 |
| optimizer | ADAM |
| loss function | Pixel-wise cross entropy loss |

**Supplemental Table 6. Autoencoder Parameters**

| Parameter | Value |
|---|---|
| batch size | 32 |
| learning rate | 0.0002 |
| optimizer | ADAM |
| dimension of the latent vector | 128 |
| dimension of the latent vector encoding texture | 4 |
| dimension of the latent vector encoding colour | 16 |
| dimension of the latent vector encoding warping | 128 |
| loss function | see [14] |

**Supplemental Table 7. Classification Performance of the *k*-nearest Neighbour Classifier Using Discriminative Phenotypic Signatures**

| Neighbour = 1 | Fold 1 | Fold 2 | Fold 3 | Fold 4 | Fold 5 | Overall |
|---|---|---|---|---|---|---|
| Precision | 1.00 | 1.00 | 1.00 | 0.88 | 0.70 | 0.91 |
| Recall | 0.60 | 1.00 | 1.00 | 0.78 | 0.78 | 0.83 |
| F1-score | 0.75 | 1.00 | 1.00 | 0.82 | 0.74 | 0.87 |
| AUC (95% CI) | 0.80 (0.64, 0.90) | 1.00 (1.00, 1.00) | 1.00 (1.00, 1.00) | 0.83 (0.78, 1.00) | 0.70 (0.47, 0.86) | 0.87 (0.80,0.92) |

| Neighbour = 3 | Fold 1 | Fold 2 | Fold 3 | Fold 4 | Fold 5 | Overall |
|---|---|---|---|---|---|---|
| Precision | 1.00 | 1.00 | 1.00 | 0.88 | 0.78 | 0.93 |
| Recall | 0.80 | 1.00 | 1.00 | 0.78 | 0.78 | 0.88 |
| F1-score | 0.89 | 1.00 | 1.00 | 0.82 | 0.78 | 0.90 |
| AUC (95% CI) | 0.89 (0.73, 1.00) | 1.00 (1.00, 1.00) | 1.00 (1.00, 1.00) | 0.90 (0.77, 1.00) | 0.82 (0.59, 0.98) | 0.93 (0.87,0.97) |

| Neighbour = 5 | Fold 1 | Fold 2 | Fold 3 | Fold 4 | Fold 5 | Overall |
|---|---|---|---|---|---|---|
| Precision | 1.00 | 1.00 | 1.00 | 0.88 | 0.80 | 0.93 |
| Recall | 0.80 | 1.00 | 1.00 | 0.78 | 0.89 | 0.90 |
| F1-score | 0.89 | 1.00 | 1.00 | 0.82 | 0.84 | 0.91 |
| AUC (95% CI) | 0.88 (0.70, 1.00) | 1.00 (1.00, 1.00) | 1.00 (1.00, 1.00) | 0.90 (0.77, 1.00) | 0.84 (0.64,0.98) | 0.93 (0.88,0.98) |

| Neighbour = 7 | Fold 1 | Fold 2 | Fold 3 | Fold 4 | Fold 5 | Overall |
|---|---|---|---|---|---|---|
| Precision | 1.00 | 1.00 | 1.00 | 0.88 | 0.80 | 0.93 |
| Recall | 0.80 | 1.00 | 1.00 | 0.78 | 0.89 | 0.90 |
| F1-score | 0.89 | 1.00 | 1.00 | 0.82 | 0.84 | 0.91 |
| AUC (95% CI) | 0.88 (0.70, 1.00) | 1.00 (1.00, 1.00) | 1.00 (1.00, 1.00) | 0.90 (0.77, 1.00) | 0.85 (0.64, 0.99) | 0.93 (0.88,0.98) |

| Neighbour = 9 | Fold 1 | Fold 2 | Fold 3 | Fold 4 | Fold 5 | Overall |
|---|---|---|---|---|---|---|
| Precision | 1.00 | 1.00 | 1.00 | 1.00 | 0.80 | 0.96 |
| Recall | 0.80 | 1.00 | 1.00 | 0.78 | 0.89 | 0.90 |
| F1-score | 0.89 | 1.00 | 1.00 | 0.88 | 0.84 | 0.92 |
| AUC (95% CI) | 0.89 (0.73, 1.00) | 1.00 (1.00, 1.00) | 1.00 (1.00, 1.00) | 0.92 (0.80, 1.00) | 0.91 (0.76, 1.00) | 0.95 (0.90, 0.99) |

AUC: area under the receiver operating characteristic curve

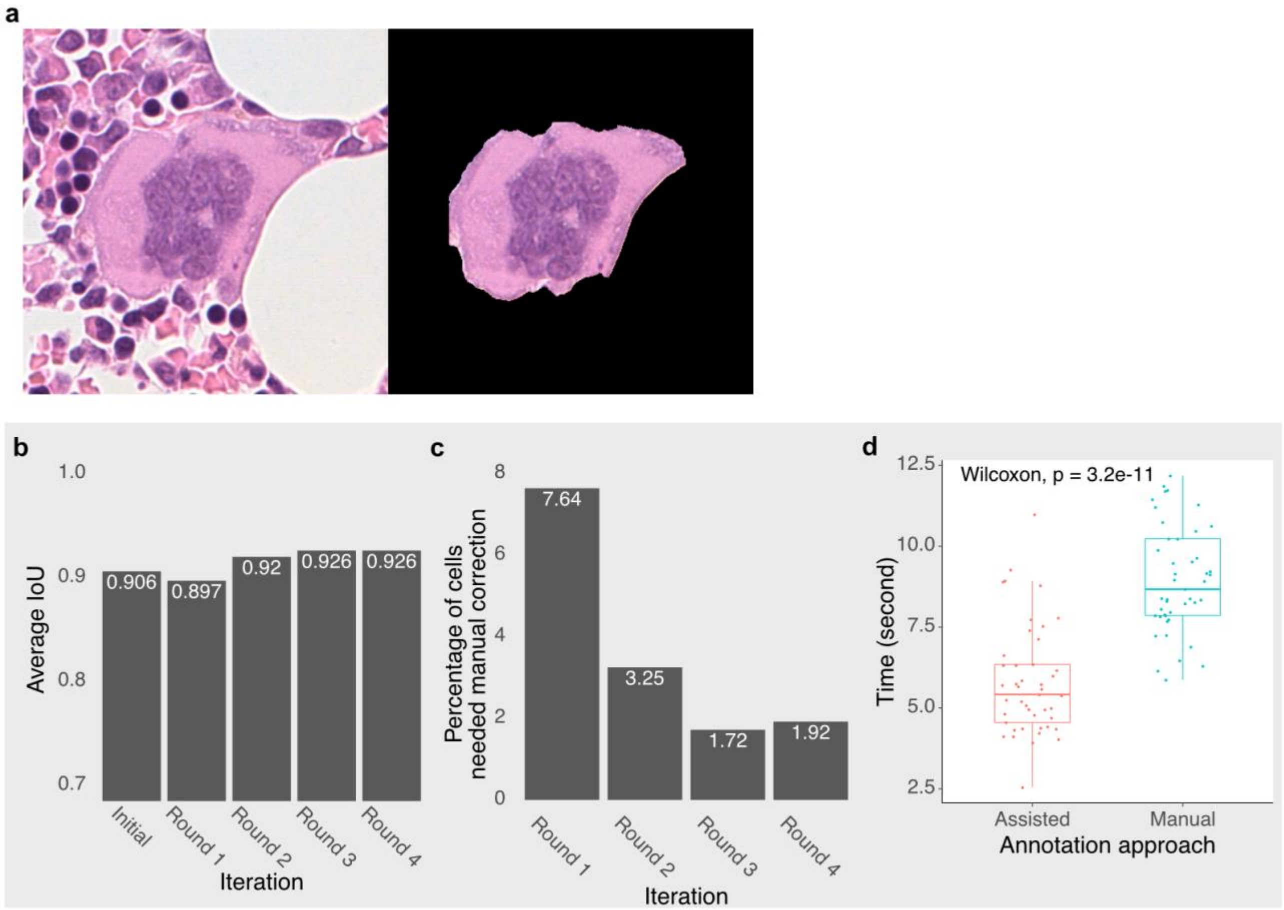


**Supplemental Figure 1: Efficiency of the Assisted Tool for Megakaryocyte Delineation.** Example of an automatically delineated megakaryocyte (**a**). The human-in-the-loop assisted annotation tool for megakaryocyte delineation achieved a high level of accuracy (IoU = 0.91) since the initial training. The performance was incrementally improved and reached IoU = 0.93 in the subsequent training rounds (**b**). The number of delineation results that needed manual correction significantly reduced in the subsequent training rounds after the initial training (**c**). The amount of time required to delineate individual megakaryocytes using our tool was significantly lower than that of manual annotation (**d**).

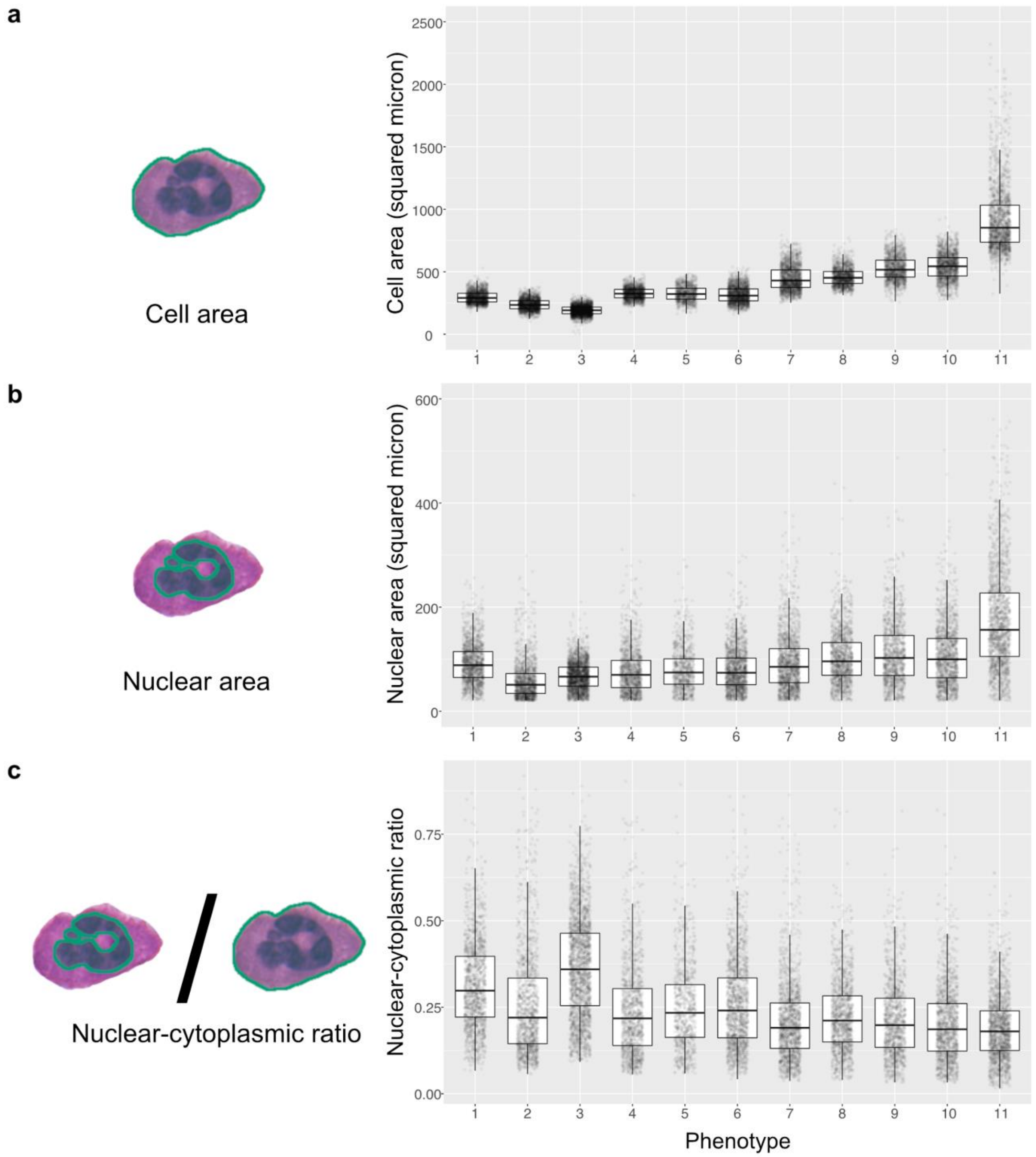
a
b
c
Cell area
Nuclear area
Nuclear-cytoplasmic ratio
Cell area (squared micron)
Nuclear area (squared micron)
Nuclear-cytoplasmic ratio
Phenotype
2500
2000
1500
1000
500
0
600
400
200
0
0.75
0.50
0.25
0.00
1
2
3
4
5
6
7
8
9
10
11

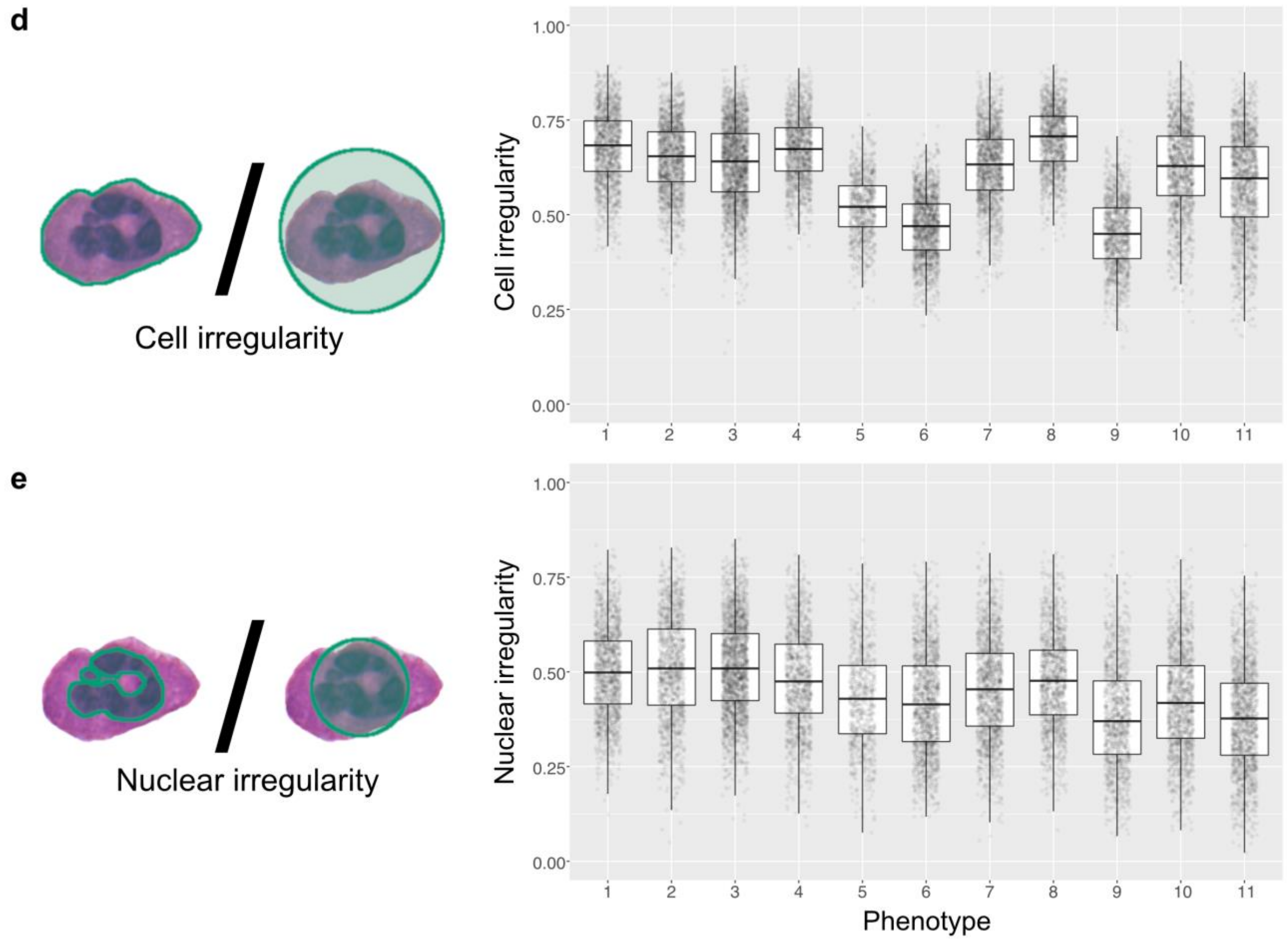


**Supplemental Figure 2. Statistics of Megakaryocyte Phenotypes.** Statistical analysis of megakaryocytes by cell area (**a**), nuclear area (**b**) and nuclear-cytoplasmic ratio (**c**). The cell irregularity was defined by the ratio of cell area to the smallest enclosing circle of the cell (**d**), with ranges between 0 and 1, where the value of 1 indicates a perfect circle (no irregularity in the shape). Nuclear irregularity was defined by the ratio of the nuclear area to the area of the smallest enclosing circle of the nucleus (**e**), with ranges between 0 and 1, where the value of 1 indicates a perfect circle (no irregularity in the shape).

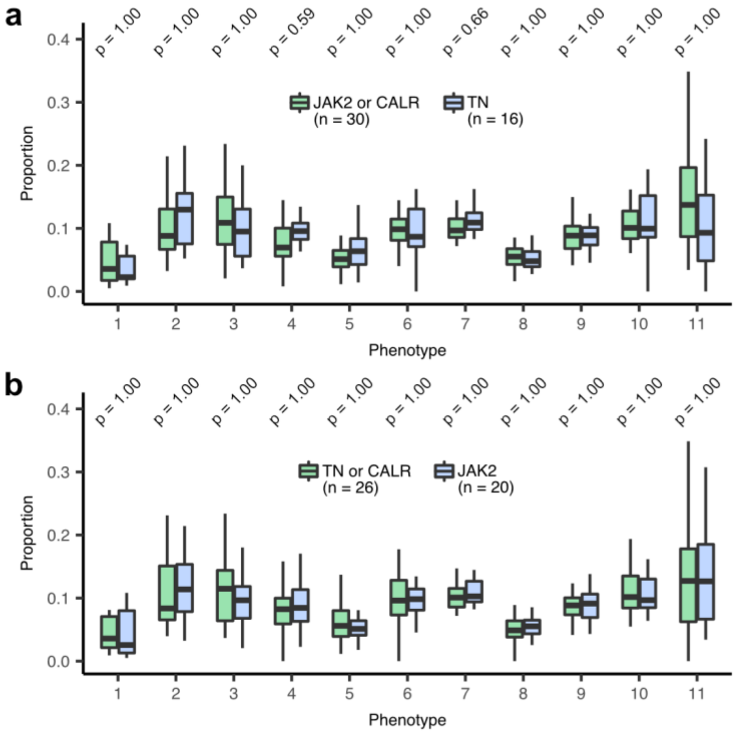


**Supplemental Figure 3: Associations Between Morphological Phenotypes and Mutational Status.** *JAK2* or *CALR* mutated versus TN ET (**a**) and TN or *CALR* versus *JAK2* mutated ET (**b**). No statistically significant result was observed. Bonferroni adjusted Wilcoxon rank-sum test, $p < 0.05$.

1. Everingham M, Van Gool L, Williams CK, Winn J, Zisserman A. The pascal visual object classes (voc) challenge. *International journal of computer vision*. 2010;88(2):303-338.
2. Alexander JB. Imgaug - python library; 2018.
3. Larsson G, Maire M, Shakhnarovich G. Learning representations for automatic colorization. Conference on Computer Vision Springer, Cham; 2016:pp. 577-593.
4. Zhang R, Isola P, Efros AA. Colorful image colorization. European conference on computer vision: Springer, Cham; 2016:649-666.
